# Spatiotemporal recruitment of inhibition and excitation in the mammalian cortex during electrical stimulation

**DOI:** 10.1101/2022.06.03.494729

**Authors:** Maria C. Dadarlat, Yujiao Jennifer Sun, Michael P. Stryker

## Abstract

Electrical stimulation has emerged as a powerful and precise treatment in which to modulate aberrant neural activity patterns common in neural dysfunction and disease; however, the physiological process involved in microstimulation is poorly understood, particularly regarding the contributions of inhibitory neurons to shaping stimulation-evoked activity. To address this issue, we used 2-photon imaging of transgenic mice to measure the widespread responses of inhibitory and excitatory neurons to electrical stimulation through a chronically-implanted cortical microelectrode. We found that increasing stimulation amplitude both raised the fraction of neurons that responded to a stimulus and increased the distance at which inhibitory and excitatory neurons were significantly modulated by stimulation; however, the lateral spread of inhibitory activity preceded that of excitatory activity. By 50 *µ*A, a significantly larger fraction of inhibitory neurons vs. excitatory neurons were modulated by stimulation. Increasing amplitude also shifted the temporal response properties of the population — towards longer-latency excitation close to the electrode tip and strong inhibition of more distant neurons. Animal behavior, specifically their locomotion patterns, strongly correlated with trial-to-trial variability in stimulation-evoked responses. We conclude that changing electrical stimulation amplitude can shift the balance of excitation to inhibition in the brain in a manner that interact with ongoing animal behavior.

## 1 Introduction

Normal brain function depends on tightly-balanced circuits of excitatory and inhibitory neurons that coordinate activity within and across brain areas to allow sensory, motor, and cognitive function. Too much or too little inhibition or excitation leads to abnormal neural (and ultimately behavioral) patterns — a problem that exists in disorders such as coma, Epilepsy, Schizophrenia, and Autism [14] amongst others. *Electrical stimulation* of the brain has emerged as a powerful and precise technique with which to modulate aberrant neural activity patterns in pharmacologically-resistant patients, supported by massive recent developments in industry that permit recording and stimulation at hundreds and even thousands of sites through chronically-implanted electrode arrays. These technological breakthroughs portend a revolution in our ability to manipulate neural activity clinically in the brains of patients by passing small electrical currents to excite or inhibit nearby neurons. However, a major obstacle in our ability to treat neural disorders using electrical stimulation is the present lack of knowledge regarding the specific stimulation patterns needed to to activate or inhibit select neural circuits in the brain.

To be able to precisely manipulate neural activity within the brain, we must answer an old and pressing question – how does electrical stimulation modulate neural activity patterns in the brain? [19]. To quote James Ranck from his 1975 review on the topic,

> The phrase ‘electrical stimulation of the lateral hypothalamus’ is a shortened version of the statement that ‘there was a stimulating electrode in the lateral hypothalamus which affected an unknown number and unknown kinds of cells at unknown locations in the vicinity of the electrode’.

We’re no longer quite at that state. The question was traditionally difficult to answer using electrophysiology, as passing stimulation current evoked a massive stimulus artifact in neural recordings which obscured evoked responses. Furthermore, the sample size of neurons collected by electrophysiology was small, and biased towards neurons with large action potentials and high firing rates. Nevertheless, data regarding the matter was steadily collected, and some coherent facts emerged. First, the prototypical response of a neuron to a single pulse or a high-frequency burst of stimulation is short-latency excitation followed by longer-lasting inhibition and, with sufficient current, a period of rebound excitation [3, 21]. Second, larger currents are generally needed to activate neurons with cell bodies farther from the electrode [23, **?**, 3, 2]; however, some distant neurons are activated at lower currents, presumably because their cell processes (dendrites or axons) pass close to the electrode tip [10].

Beyond establishing these general rules, we do not yet understand the neural circuits that mediate responses evoked by electrical stimulation, particularly the role of inhibitory neurons in shaping stimulation-evoked activity patterns. The spatiotemporal patterns of neural population activities is determined by multiple factors, shaped by the dynamic interactions between excitatory and inhibitory signals [25]. Inhibitory networks shape the spatial spread and temporal response of both natural and stimulation-evoked neural activity in the the cortex [9, 2]. However, we don’t yet know how spatial and temporal activation of inhibitory neurons shape evoked neural responses.

One direct approach towards recording the activity of inhibitory neurons is use to genetic techniques to label distinct neuron classes, and to record neural activity from each type of neuron. This can be most easily achieved by optical imaging. Laser scanning 2-photon imaging of calcium signals of neuronal activity has two advantages over electrophysiology and one major disadvantage. The disadvantage is that it is slow compared to electrophysiology. Calcium levels have slow decay times and scanning permits only low sampling rates that prohibit measurement of the precise timing of neural responses. However, the advantages of imaging are significant. Foremost is the lack of stimulation artifact, meaning that directly-activated neural responses can be recorded. Furthermore, powerful genetic tools permit labeling of distinct cell classes, which is needed to understand the circuits involved in shaping stimulation-evoked neural responses. Finally, the population of neurons that can be recorded simultaneously within imaging reaches thousands, permitting a comprehensive, detailed picture to be formed of the spread of excitation and inhibition across the cortex following stimulation.

The use of 2-photon imaging has been critical for expanding our understanding of how electrical currents can shape neural responses patterns. We know, for example, that neurons activated by electrical stimulation are sensitive to the temporal pattern of stimulation, and spatiotemporal activation of neural populations is achieved by changing the frequency and precise timing of delivered stimulation pulses [5, 12, 7, 22]. But maybe the first study using 2-photon imaging to examine stimulation-evoked neural activity was the most dramatic in how it shifted our perception of how electrical stimulation works. In 2009, Histed and colleagues reported that, rather than activating neurons that lay within a compact sphere surrounding the electrode tip, electrical stimulation directly activated a sparse and distributed subset of neurons whose axons were thought to pass within tens of microns from the electrode tip. Why the discrepancy? To address this issue, we used 2-photon imaging to record activation patterns of genetically-labeled inhibitory and excitatory neurons in response to increasing stimulation currents passed through a Pt/Ir electrode that was chronically-implanted in mouse visual cortex while animals were head-fixed but awake and free to run or rest on a spherical treadmill.

## 2 Methods

### Surgery

All procedures were conducted in accordance with the ethical guidelines of the National Institutes of Health and were approved by the Institutional Animal Care and Use Committee at the University of California, San Francisco. The surgical procedures used in these experiments have been described previously [5] and are related briefly below. The overall experimental setup is shown in Figure 1.

**Figure 1:**
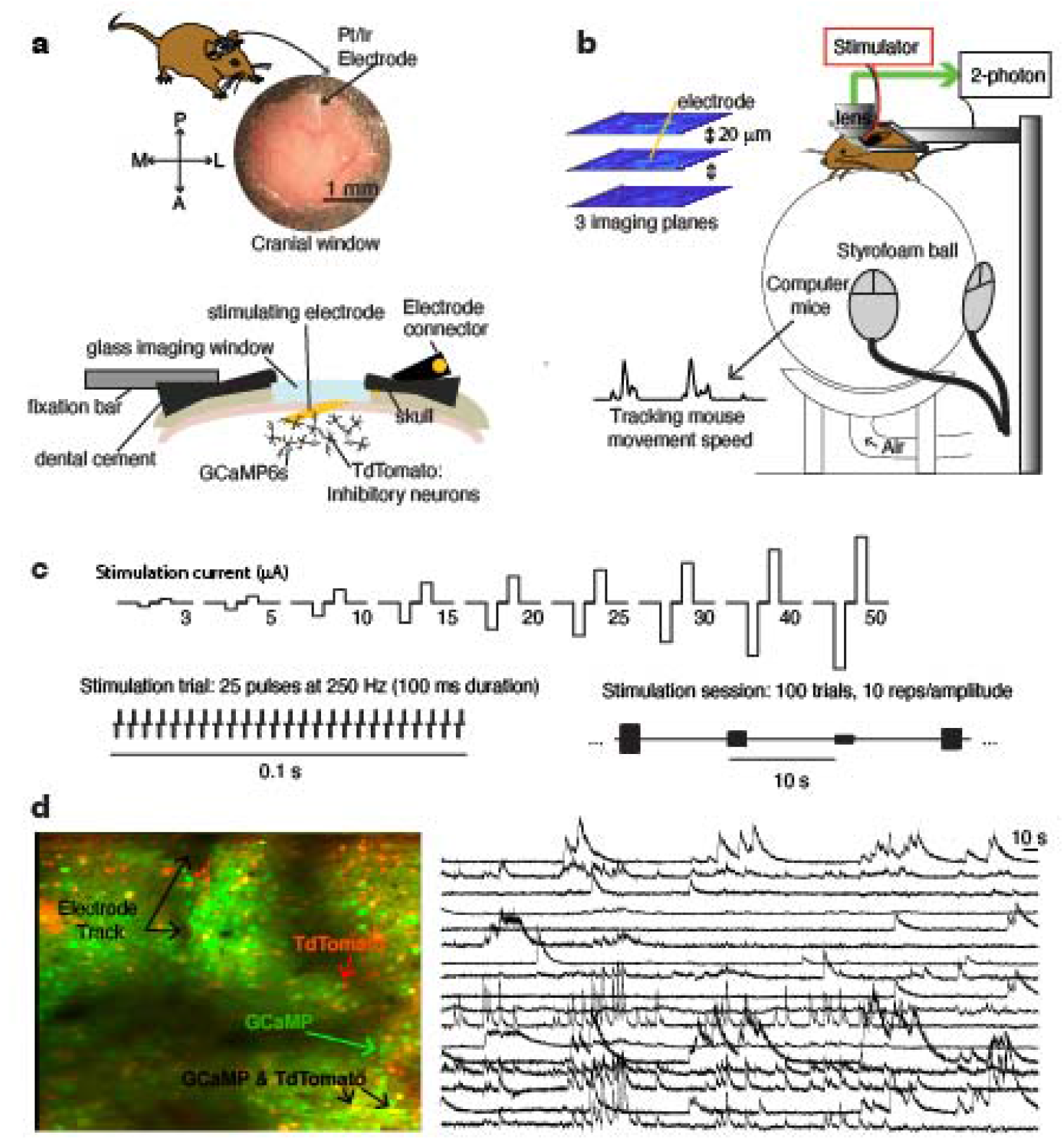
Neural recording and stimulation in chronically-implanted awake mice. **a)**. Experimental setup: awake mice are head-fixed and free to run or rest on a spherical treadmill. Three cortical planes were imaged during each imaging session. **b)**. A single Pt/Ir stimulating electrode was chronically implanted in mouse primary visual cortex beneath a glass imaging window. **c)**. Electrical stimulation consisted of 25 biphasic cathode-leading pulses delivered at 250 Hz. Stimulation was delivered once every ten seconds at one of nine possible current amplitudes: 3, 5, 10, 15, 20, 25, 30, 40, or 50 *µ*A. Ten stimuli were delivered at each amplitude. **d)**. Imaging was conducted in red and green channels; sample imaging plane at 2x to see detail (experimental sessions were conducted at 1x). At right are sample time-varying signals reporting changes in fluorescent intensity from a subset of regions of interest.

Inhibitory neurons in mice were labeled with red fluorescent markers by crossing transgenic mouse lines expressing Gad2-Cre with those expressing floxed-TdTomato (Jackson labs strains 010802 and 007914). We studied five mice of ages 3–6 months (both male and female) were used in this study. Each mouse underwent a total of three separate surgical procedures in order to implant 1) a titanium headplate, 2) a platinum/iridium (Pt/Ir) electrode (0.1 MΩ, Microprobes for Life Sciences) and a glass cranial window, and 3) a two-pin receptacle connector and ground wire. Just prior to implantation of the stimulating electrode, mice were injected with 100 nL of AAV virus expressing GCaMP6s into three locations within primary visual cortex (AAV1.Syn.GCaMP6f.WPRE.SV40; Addgene 100837-AAV1) at two depths (150 and 300 *µ*m) per location thereby transfecting both excitatory and inhbitory neurons with a fast fluorescent indicator of calcium concentration. For each procedure, animals were anesthetized to areflexia using a mixture of Ketamine and Xylazine (100 mg/kg and 5 mg/kg, IP).

The surgery to implant a titanium headplate was previously described in detail [4, 15]. The headplates used here were modified from the original design by removing the rear third, leaving a semicircular portion with lateral flanges. We implanted the Pt/Ir stimulating electrode and glass cranial window between three to seven days after the headplate surgery. Implantation of the glass cranial window closely follows well-established experimental protocols [8, 6]. The stimulating electrode was implanted after a craniotomy was made to expose the cortical surface. Following drilling of the skull to the point that it could be removed but prior to its removal, a small ramp was shaved in the skull just anterior to the craniotomy. This ramp allowed us to implant the stimulating electrode such that it entered the cortex at the very edge of the cranitomoy – a detail that was critical for successfully securing the glass cranial window and ensuring a long-lasting chronic implant.

The Pt/Ir electrode was inserted at a 7*°* or 11*°* angle to a depth of 200 µm below the cortical surface using a digital micromanipulator (Sutter MP-285). Next, a layer of sterile Vaseline was placed along the posterior of the craniotomy behind which a layer of cyanoacrylate mixed with dental acrylic was added to fix the electrode in place. After the electrode was secure and the Vaseline removed, a 3 mm glass window was placed over the craniotomy. The edges of the window were fixed in place using cyanoacrylate, and were further stabilized by the application of dental acrylic. The electrode was clipped with wire cutters at its exit from the dental cement, leaving only an edge exposed.

After allowing the animal to recover for several days, a two-pin receptacle connector and ground wire were implanted. One pin of the connector was electrically connected to the end of implanted electrode using a conductive silver paint. The connection was insulated using cyanoacrylate, and the rest of the connector was firmly attached to the rear of the titanium headplate using more cyanoacrylate. To complete the circuit, a small hole was drilled through the skull over the contralateral frontal lobe. The stripped end of a chloridized silver wire was inserted beneath the skull (making contact with cerebral spinal fluid but not piercing the brain) and the opening was covered with a biocompatible silicone elastomer. Finally, cyanoacrylate was applied over the silicone elastomer and over any loose portion of the ground wire to provide stability and electrical insulation.

### Electrical Stimulation

The goal of the experimental protocol was to understand how electrical stimulation of a range of amplitudes modulates activity in surrounding neurons and to isolate differences in its effect on excitatory and inhibitory neurons. The stimulation protocol consisted of delivering a constant-current electrical stimulus of nine different intensities once every ten seconds, for a total of ninety stimuli during the recording session (fig. 1c). The amplitude of each stimulus was chosen from an array of possible values (3, 5, 10, 15, 20, 25, 30, 40, or 50 *µ*A) in a psuedo-random fashion, for a total of ten repetitions per stimulus amplitude across the experimental session. Stimulation frequency was held fixed at 250 Hz. Each stimulus consisted of a train of twenty-five biphasic, cathode-leading pulses for a total stimulation duration of 100 ms. Biphasic pulses were composed of a 200 *µ*s cathodal pulse, a 200 *µ*s inter-pulse interval, and a 200 *µ*s anodal pulse.

### Two-photon calcium imaging

Calcium responses of specific cell types and processes were acquired using a resonant-scanning two photon microscope (Neurolabware) controlled by Scanbox acquisition software. A Coherent Chameleon Ultra II laser was used for GCaMP excitation at a wavelength of 920 nm. Emission light was filtered by emission filters (525/70 and 610/75 nm) and measured by 2 independent photomultiplier tubes (Hamamatsu R3896). The objective used was a 16 water-immersion lens (Nikon, 0.8 numerical aperture (NA), 3 mm working distance). Image sequences were captured at 15.5 Hz at a depth of 100–300 *µ*um below the pia. All recording was done with awake, head-fixed mice that were free to walk or run on a spherical treadmill.

Neural responses were imaged in three planes spaced 20 *µ*m apart in depth, covering an area of approximately one *µ*m^2^ (1136 *µ*m by 1083 *µ*m). Each plane imaged at 5.16 Hz (15.5 Hz divided by three imaging planes). In four of the mice, the first imaging plane was placed the depth of the stimulating electrode (assessed by visual inspection or automated z-stack imaging) and the remaining two planes were 20 and 40 *µ*m deeper. In one of the mice with a deeper implanted electrode we instead placed imaging planes at the electrode tip and 20 *µ*m above and below the electrode tip.

During imaging, awake mice were head-fixed and placed upon a Styrofoam sphere that floated on a bed of air. Upon the sphere, mice could freely run or stay still, according to the inclination of each (fig. 1b); [6]). All neural recordings took place in a dark room and the animals were isolated from the experimenter in an enclosure formed by black laser safety fabric. Before recording, mice underwent at least 5 days of training to acclimate to the treadmill and head-fixation. Treadmill movement was captured by a Dalsa Genie M1280 camera synchronized to microscope scanning.

### Analysis

#### Registration, cell detection, and neuropil correction

Image registration, region of interest (ROI) detection, neuropil correction and deconvolution of the two-photon imaging data were carried out using the Suite2p pipeline ([17]). The data were further curated through manual cell detection, differentiating between cell and non-cell ROIs using the suite2p GUI. Each ROI identified by suite2p was manually analyzed to determine whether suite2p’s classification of cell or non-cell was correct. The classifier was trained by considering the calcium traces, shape, compactness, aspect ratio and position of the ROIs. In the post-stimulation sessions, ROIs were considered as cells if: 1. they had calcium traces considerably different from the surrounding neuropil activity, and 2. a round, compact shape of a size large enough to represent a cell body rather than the cross section of a dendrite. We kept all ROIs that were assigned by Suite2p at least a probability of 0.5 of being a neuron. All subsequent analyses were conducted in Matlab using custom code.

Electrical stimulation induced a large, synchronized deviation of the neuropil signal, which may bias the responses recorded in “single-neuron” ROIs. Therefore, to isolate a single neuron’s activity across the imaging session, we subtracted 0.8 times the neuropil signal surrounding each cell (neuropil was assumed to be an annulus surrounding each ROI). Note that this value is larger than that conventionally used (e.g. [24]) to account for the high level of neuropil synchrony during electrical stimulation.

#### Identification of inhibitory neurons

Red cells (genetically labeled inhibitory neurons) were found by detecting ROIs from the red channel, after the correction of the green channel dynamics using the linear fitting. If the corrected red ROI signals were significantly greater than the local average (2 time greater than the standard deviation), the cells were classified as inhibitory neurons.

#### Radial distance from electrode tip

In order to describe stimulation-evoked activity patterns as a function of radial distance from the electrode tip, each neuron’s relative position was calculated as follows. The position of the electrode tip was found by visually inspecting the max projection image across the aligned imaging movie data. Within each imaging plane, a neuron’s position was said to be at the mean of the × and y pixels assigned to that ROI, which were then converted into *µ*m. We found radial distance for each neuron, r, by including information about the imaging plane in which each ROI was located: 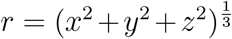 with *x* and *y* are the mean position of a neuron within the imaging plane (in mm) and *z* is the distance between the plane of the electrode tip and that of the ROI (0, 20, or 40 *µ*m). Once the relative position of each cell was found, we binned all distances into bins of width 100 *µ*m, ranging from 0 to 500 *µ*m.

#### Significant modulation of neural activity

Neural responses to stimulation were characterized as the change in fluorescent intensity of the cell body for 20 samples relative to its value at stimulation onset. This data spanned approximately 4 seconds at the 5.17 Hz sampling rate per imaging plane, with sample 1 being the time at which stimulation occurred. The responses were split into two time periods: an early stage (0-800 ms after stimulation) and a late stage (800 - 3800 ms after stimulation). Then early responses were said to be the average of the neural fluorescence in the early stage and late responses were said to be the average of neural fluorescence during the late stage. To determine significant modulation by electrical stimulation, we compared early and late stage responses for the ten trials at each amplitude with the neuron’s single-trial baseline activity averaged over the period between from four to one second prior to stimulation onset. Significant deviation from baseline was tested for the early and late stage separately using a Wilcoxon rank-sum test, taking p ≤ 0.02 as a threshold. No corrections were made for multiple comparisons; instead, further analysis takes this level of activation (one out of every 50 neurons considered) to be equal to chance.

#### Amplitude, variance, and Fano Factor of evoked neural responses

The amplitude of an evoked neural response was calculated in one of two ways. The first method, used in fig. 2, simply takes the maximum value of a neuron’s average response (a four second trace of fluorescence as a function of time, beginning at the time of stimulation) to each stimulation amplitude (averaged across ten trials delivered at each current amplitude). To account for neurons that were predominantly inhibited by stimulation, we also find the minimum of the trial-averaged response trace, and take as the modulation amplitude the maximum of the absolute value of the minimum of the response and the maximum of the response: max(|*y*_*min*_|, *y*_*max*_), where y is the time-varying trace of the averaged response for a neuron. The second method, used in fig. 5, takes into account the temporal response of each neuron (see below description of neural response profiles and fig. 4) when calculating response amplitude. In this case, the response amplitude was taken to be the average value of the response stage in which the neuron was significantly modulated. Because this analysis splits apart inhibitory from excitatory responses, we did not take the absolute vale of inhibitory responses. If the neuron was significantly modulated during both stages, we took the response amplitude to be the maximum of the absolute value of the average activity evoked in each stage (to account for inhibitory responses).

**Figure 2:**
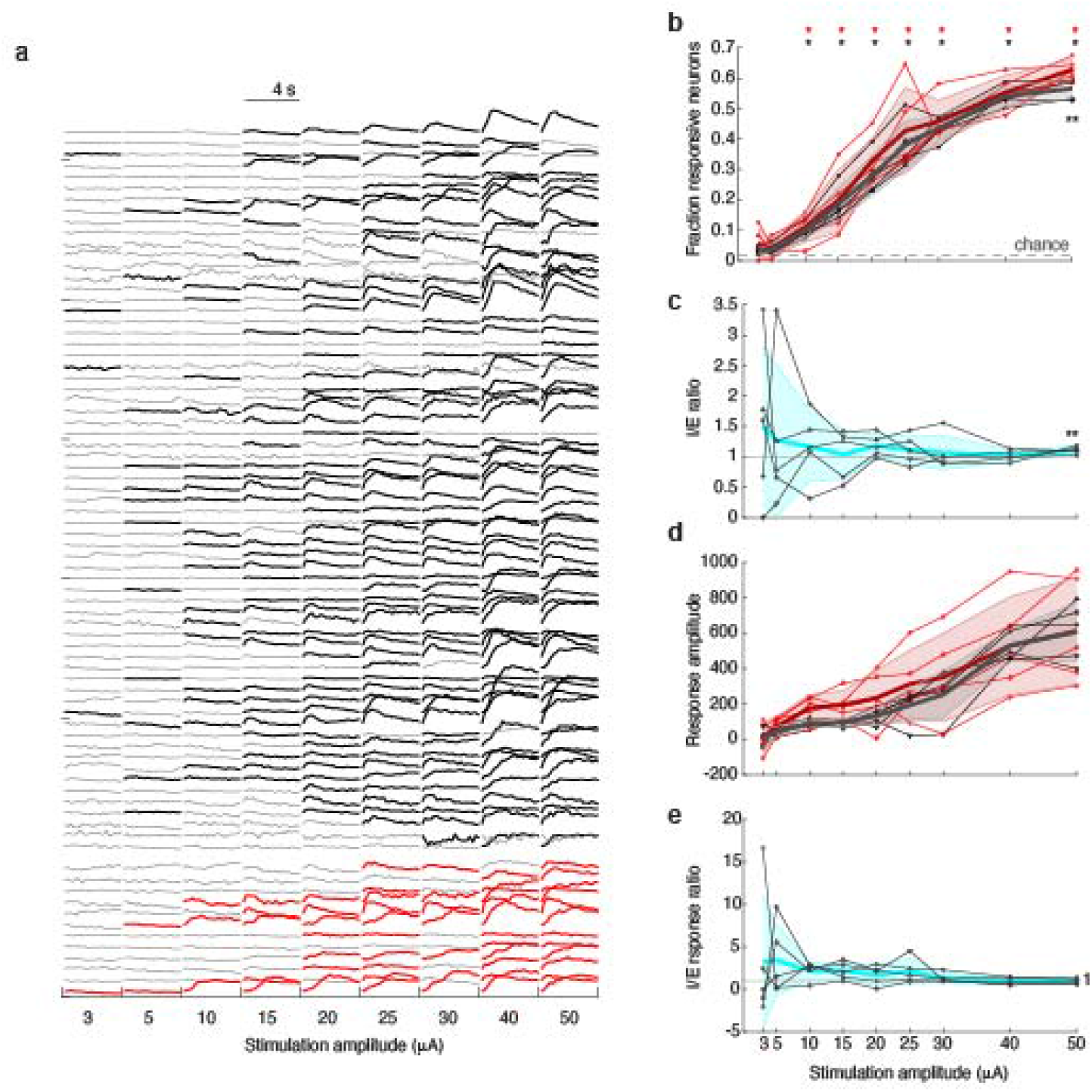
Neural activity evoked by electrical stimulation. **a)**. Sample evoked responses from neurons with increasing stimulation intensity, averaged across 10 trials. Each row is a single neuron. Significant responses in black (excitatory neurons) or red (inhibitory neurons) (p≤ 0.02; Wilcoxson rank-sum test). **b)**. Fraction of neurons significantly modulated by stimulation as a function of current amplitude. Red is inhibitory; black excitatory. Bold lines show averaged data; thin lines show single-mouse data. Shaded error bars are STD. Dashed gray line shows chance level. Significance above chance is indicated by *\** at top of the panel. Significant difference between inhibitory and excitatory traces are shown by *\* \** (p ≤0.01, t-test) **c)**. Ratio of significantly responding inhibitory cells to excitatory cells. Bold lines show averaged data; thin lines show single-mouse data. Shaded error bars are STD. Gray line is at one, where the fraction of evoked responses are equal for inhibitory and excitatory neurons. Significant difference between the two (deviation of the ratio from one) is indicated by the *\* \**symbols above the data (p≤ 0.01, t-test). **d)**. Average amplitude of significant evoked neural responses to stimulation. Colors and lines as in (b). **e)**. Ratio of inhibition evoked responses to excitatory evoked responses. Colors and lines as in (c).

**Figure 3:**
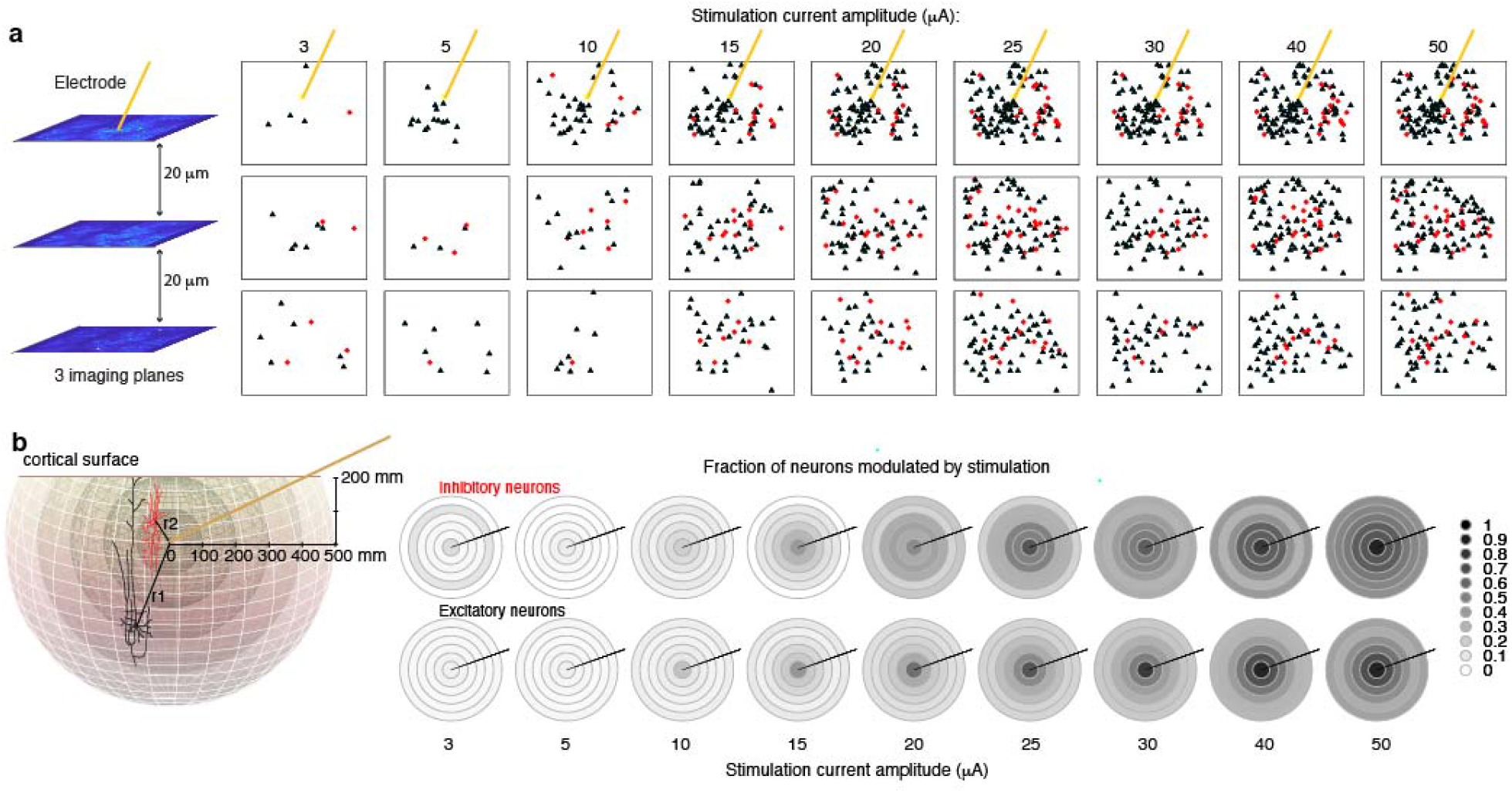
Neural responses to stimulation depend on the distance between the cell body and the stimulating electrode tip. **a)**. Neural activation as a function of current amplitude within the three imaging planes. Significant responses marked in black (excitatory) and red (inhibitory). **b)**. Radial spread of stimulation-based modulation, from 100 microns. Colors indicated fraction of neurons modulated by stimulation.

**Figure 4:**
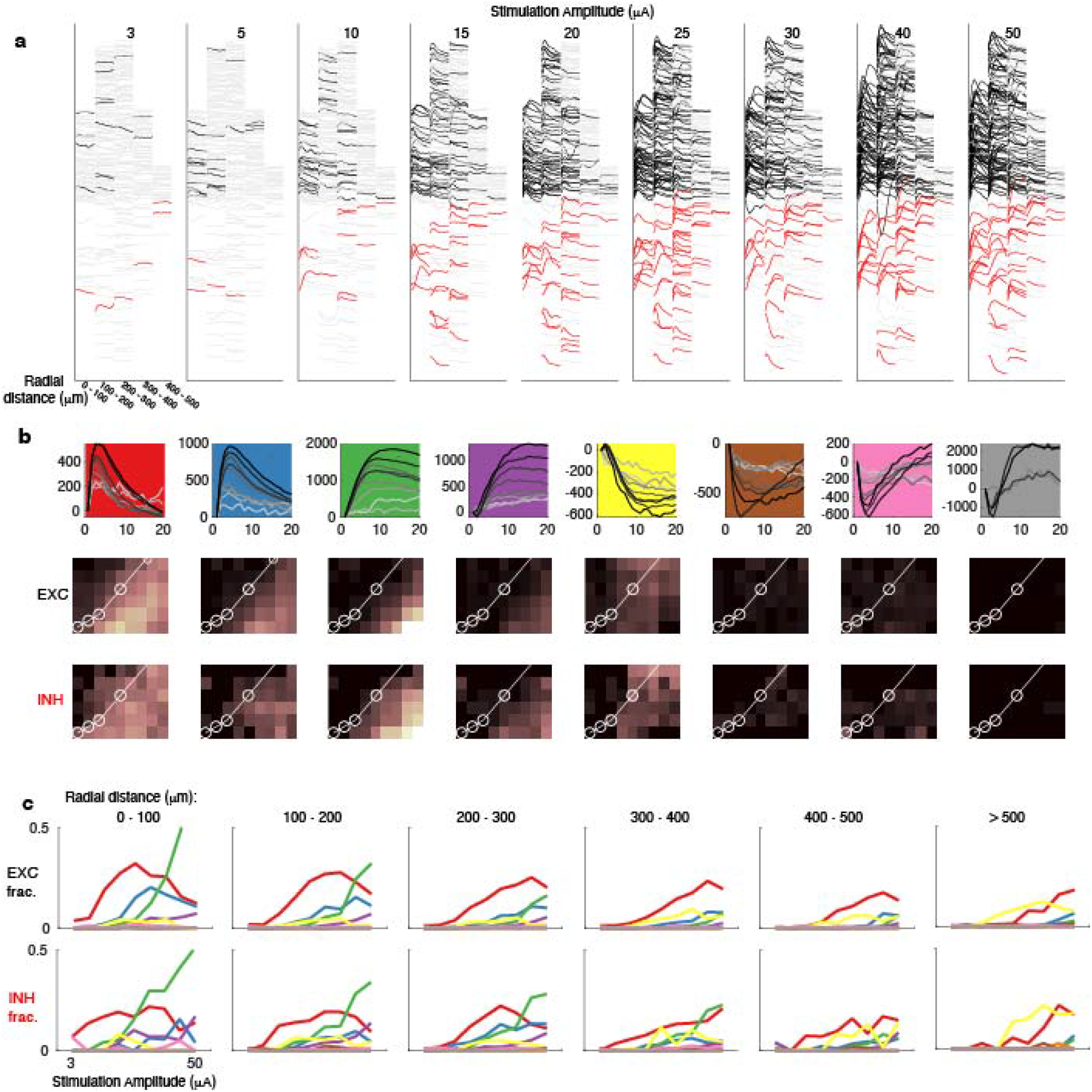
Spatio-temporal neural response patterns to electrical stimulation. **a)**. Samples responses to electrical stimulation from mouse 1, separated by radial distance from stimulating electrode (columns within each panel). Color traces indicate significant modulation of excitatory (black) and inhibitory (red) neurons. **b)**. Top panels: five dominant response profiles for neurons modulated by electrical stimulation. Bottom panels: spatial distribution of each type of response as a function of stimulation amplitude. White lines show data of expected spatial activation patterns from [19]. **c)**. Fraction of each response profile as a function of stimulation amplitude, separated into panels by average radial distance from stimulation electrode. Color as in (d).

**Figure 5:**
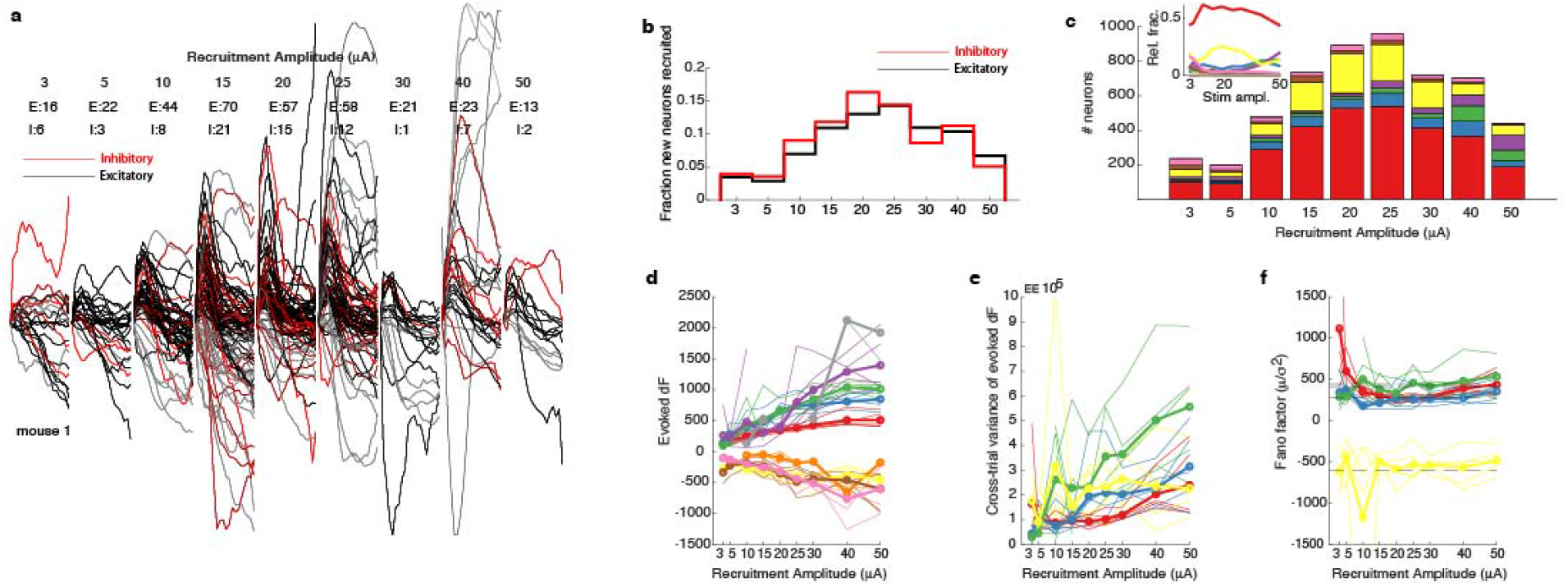
Recruitment amplitude of neurons. **a)**. Sample responses of newly-recruited neurons from mouse 1. **b)**. Distribution of recruitment amplitudes for inhibitory and excitatory neurons across all experimental animals. **c)**. Response types of neurons at each recruitment amplitude. Colors are as in fig. 3d. Inset shows relative fraction of each response type. **d)**. Average amplitude of evoked response for neurons recruited at each stimulation amplitude, separated by response type (colors as in fig. **??**). **e)**. Variability of evoked response amplitudes across single trials for neurons recruited at each stimulation amplitude, separated by response type (colors as in fig. 3). **f)**. Fano factor 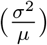 of response amplitudes for neurons recruited at each stimulation amplitude, separated by response type (colors as in fig. 3). Horizontal line shows mean across all stimulation amplitudes for each response type.

The variance of the response amplitude is calculated by taking into account trial-to-trial fluctuations. For this analysis, single-trial amplitudes were first calculated as described above, and then the standard deviation was measured across the ten trials of a single amplitude. The fano factor of the response amplitudes is calculated as the variance of the response amplitude divided by the mean, 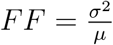.

#### I/E ratio

We measured ratio of modulation of inhibitory to excitatory neurons by electrical stimulation (I/E ratio) for both the fraction of neurons modulated by electrical stimulation as well as for the response amplitude of these neurons. The I/E ratio was found by simply dividing the average fraction of inhibitory neurons activated (or average response amplitude across all inhibitory neurons) per stimulation amplitude by that for excitatory neurons.

#### Establishing neural response profiles

Based on prior literature, we know that neurons can be either excited or inhibited or both by electrical stimulation at varying time lags relative to stimulation onset. In order to categorize the temporal responses of neurons and to catalog their diversity, we considered nine possible combination of significant response patterns: 1. Significant excitation only in stage 1 (2-5 samples after stimulation, which would be a fast, low-latency response 2. Significant excitation in both stages with a maximum response in stage 1 (low-latency response with slower offset) 3. Significant excitation in both stages with a maximum response in stage 2 (longer-latency peak but early onset) 4. Significant excitation only in stage 2 (long-latency response) 5. Significant excitation in stage 1 followed by significant inhibition in stage 2 (short-latency excitation followed by long-latency inhibition) 6. Significant inhibition only in stage 2 (long-latency inhibition) 7. Significant inhibition in both stages (fast-onset long-lasting inhibition) 8. Significant inhibition in stage 1 (fast-onset short-lasting inhibition) 9. Significant inhibition in stage 1 followed by significant excitation in stage 2 (fast-onset inhibition with rebound excitation).

#### Recruitment amplitude

We define neural recruitment amplitude to be the amplitude of stimulus current to which a neuron first has a significant response (found as described above). To find the recruitment amplitude for all neurons, we first created a binary significance matrix that described whether or not each neuron was significantly modulated by electrical stimulation at each of the nine stimulation current amplitudes we tested. Using this matrix, we found for each neuron the first current amplitude at which a neuron was significantly modulated by electrical stimulation and called that the neuron’s recruitment amplitude. No further restrictions were placed on the array of responses for this analysis, so we include neurons that are significantly modulated by low stimulation amplitudes but that stop being modulated by higher amplitudes.

#### Behavioral modulation of stimulation-evoked neural activity

We modeled the response of a single neuron to electrical stimulation as a linear function of the movement speed of the mouse and the amplitude of stimulus current: *n*_*i*_ = *b*_0_ + *b*_1_ *v*_*i*_ + *b*_2_ *s*_*i*_ + *ϵ*_*i*_, where *ϵ* is assumed to be iid and drawn from a standard normal distribution, *b* is a baseline offset, *b*_1_ is the weight given to *v*_*i*_ – the average speed of the mouse during trial *i*, and *b*_2_ is the weight given to *s*_*i*_ – the stimulation current amplitude for trial *i*. For this analysis, a neural response for a trial was taken to be the average value across samples 1-4 following stimulation (a period of time approximately equal to 0.75 s). Per-trial movement speed was calculated as the average speed during the 20 samples following stimulation, which is the time-course across which we saw significant modulation in neural activity patterns following stimulation. Next, we calculated *R*^2^ for each model fit to estimate how well the linear relationship captured the responses of each neuron.

## 3 Results

Altogether, we recorded neural activity from 6698 neurons across five mice: 6162 excitatory neurons and 532 inhibitory neurons. Figure 1 shows the experimental procedure and a typical image and response.

### High stimulation intensities recruit large fractions of neurons

In accordance with previous work [3], increasing the strength of the current used for stimulation progressively recruited larger fractions of both excitatory and inhibitory neurons (fig. 2a,b). A neuron’s activity pattern was considered to be significantly modulated by electrical stimulation if it deviated from baseline in one of two stages: directly after stimulation (one second following stimulation) or if it had a longer-latency deviation (during the following three-second period). Significance was tested using a Wilcoxon rank-sum test to compare between the single-trial activity during baseline prior to stimulation with the single-trial activity during each stage following stimulation (see Methods for more detail). We used p ≤0.02 as a threshold for significance for single neurons, which would lead to a false-positive rate of 2 neurons of every 100 tested. Using this value as an estimate of chance, we found that the threshold stimulus current for many inhibitory and excitatory neurons was between 5 and 10 *µ*A (fig. 2b). Additional inhibitory and excitatory neurons were recruited at approximately equal rates until 50 *µ*A stimulation, where inhibitory recruitment significantly exceeded excitatory recruitment (p ≤ 0.01, t-test; fig. 1b,c), reaching a ratio of 1.1 inhibition to excitation.

### The amplitude of evoked responses scales with stimulation current amplitude

Next, we tested whether increasing stimulation current not only recruited more neurons, but also changed the amplitude of evoked responses for single neurons. We calculated the evoked response of a neuron to stimulation as the average deviation of the flouresence trace from baseline during the time period that neuron was significantly modulated (during the early or late stage). If the neuron was significantly modulated during both time ranges, the average was taken for the time range that had the largest deviation from baseline. We found that, in general, excitatory and inhibitory neurons had statistically similar responses across all stimulation current amplitudes (fig. 2d,e). There was a trend for higher-amplitude responses from inhibitory neurons just past threshold activation levels (10 - 20 *µ*A; fig. 2d,e) but the difference was not significant.

### Radial distribution of stimulation-evoked neural activity

A major objective of our study is to relate the spread of excitation and inhibition with the amplitude of stimulation current. When plotting the spatial distribution of significantly-modulated neurons, a pattern emerges; although neural recruitment at low stimulation amplitudes is sparse and widely distributed [10], increasing stimulation amplitude raises the probability that distant neurons are significantly modulated (fig. 3a; data from mouse 1). To quantify these observations, we first calculated the radial distance between the electrode tip and the location of each neural cell body within our three imaging planes (fig. 3b left panel), pooling data from all five mice. Neurons were then separated by type (excitatory or inhibitory) and radial distance was binned by 100 *µ*m to divide distance into size groups, the first of which included neurons falling within 100 *µ*m of the electrode tip and the last of which grouped all neurons beyond 500 *µ*m from the electrode tip. Using the binned data, the probability that neurons at a certain distance from the electrode would be activated by each stimulation amplitude was calculated. We found that the probability of neurons within each bin being activated grew with stimulation amplitude; however, the lateral spread of inhibition exceeded that of excitation at each stage (fig. 3b). Thus, our data support the hypothesis that inhibitory activity curtails the lateral spread of excitation elicited by electrical stimulation. At higher stimulus amplitudes responsive neurons were farther and to the edges of the region imaged. Therefore we we cannot put an upper bound on the spread of activity.

### Spatial and temporal response properties of single neurons change with stimulation current strength

So far we have only concerned ourselves with whether or not a neuron changes its activity pattern after stimulation while completely ignoring the details of that change. However, neurons exhibit a diversity of response profiles to electrical stimulation (fig. 4), from fast excitation or inhibition to longer-latency modulation and even a mixture of the two across time. We set out to characterize these responses and hypothesized nine possible response profiles (see Methods - Establishing response profiles) of which we readily found eight (fig. 4b). Each response profile was found to a different degree in inhibitory versus excitatory cells and had a distinct spatial distribution across stimulation amplitudes (fig. 4b,c). In general, low-latency short-lasting responses dominated close to the electrode up to 25 *µ*A. We hypothesize that these are neurons whose axons or cell bodies are directly activated by electrical stimulation, which would lead to a low-latency response. However, with increasing stimulation amplitude, the dominant response pattern becomes a longer-latency high-amplitude activation. In contrast to short distances, at longer radial distances (most prominently beyond 500 *µ*m), neurons, particularly excitatory neurons, are primarily inhibited by electrical stimulation. At these longer distances, increasing current strength no longer leads to the long-latency excitation patterned that dominates short radial distances. Instead, increasing amplitude recruits more of the putative “directly-activated” neurons. At all distances the amount of inhibition slowly rises until around 30 *µ*A after which it mostly falls.

### Recruitment Amplitude

As is clear in fig. 2a, neurons have different recruitment amplitudes, which we define to be the stimulus current at which a neuron is first significantly modulated by electrical stimulation. So far, we have only considered neural recruitment in the aggregate – coarsely lumping together neurons with different recruitment amplitudes. In order to gain a finer view of the spatial distribution and response profile of neurons to electrical stimulation, we further divided neurons by recruitment amplitude. The fraction of additional neurons significantly modulated by electrical stimulation grew with stimulation current amplitude and peaked at 20 *µ*A for inhibitory neurons (recruiting a newly-responding fraction of 0.16 of all inhibitory neurons) and at 25 *µ*A for excitatory neurons (recruiting a newly-responding fraction of 0.14 of all excitatory neurons; fig. 5a,b). The distribution of response profiles changes as a function of recruitment amplitude (fig. 5c), but the dominant response (ranging from 43 % to 61 %) at all values is that of a fast-onset excitatory response onset followed by a fast decay. The prevalence of newly-recruited neurons with higher stimulus amplitudes that are primarily *inhibited* by stimulation follows a similar trajectory – gaining in number with increasing amplitude and peaking at 225 neurons during 20 *µ*A stimulation, at which point it makes up 25% off all responses. Unlike our previous analysis, the distribution of response types for newly-recruited neurons was independent of radial distance from the electrode tip (p = 0.174, 1-way ANOVA).

### Trial-to-trial variability in neural responses to stimulation

We found that neural responses to electrical stimulation were variable from trial-to-trial both in the magnitude and time-course of the response (fig. 6a,b). During natural visual processing, neural responses to visual stimuli are strongly modulated by fluctuating behavioral states including alertness and locomotion [15]. Similarly, we observed strong fluctuations in neural activity patterns evoked by electrical stimulation across the population of neurons we recorded, particularly among the most active neurons (fig. 6b). Organizing the single-neuron responses by stimulation amplitude, we can see that behavioral or state fluctuations may contribute to high variability in the amplitude of evoked responses by a single stimulation current value (fig. 6c). Despite these high levels of variability in neural responses across trials, the *average* response across the neural population increases with increasing stimulation amplitude for all of the mice in which we recorded (fig. 6d).

**Figure 6:**
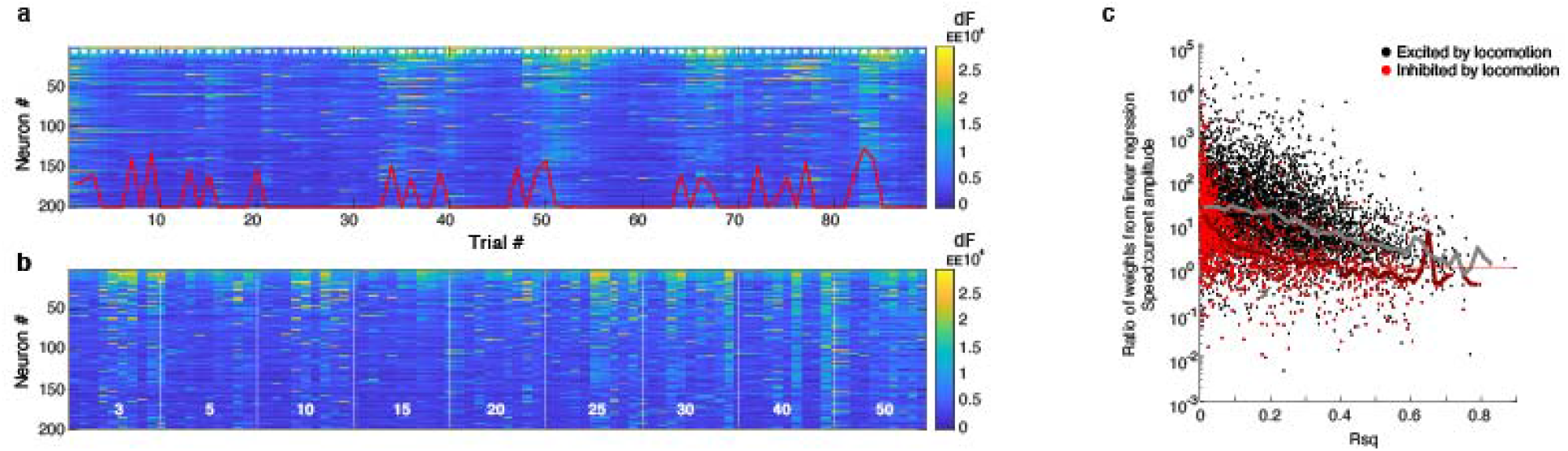
Behavioral modulation of stimulation-evoked neural activity. **a**. Average post-stimulation response of the 200 neurons with the highest average response rate across all eighty-nine stimulation trials. Red traces shows mouse movement speed for each trial; values are scaled for clarity. White numbers show stimulation current amplitude for each trial. **b**. Same data as in *a* re-arranged to group stimulation current amplitude. **c**. Ratio of weights from linear regression for single-neuron responses as a function of running speed and current stimulation amplitude. Neurons were both excited (black points) and inhibited (red points) by locomotion. Median values for each group shown by lines.

To clarify the contribution of fluctuating behavioral states to stimulation-evoked neural activity patterns, we decided to see if locomotion speed (a behavioral measures tracked throughout the experiments; fig. 1b) was the key driver of variability in the responses of individual neurons to stimulation. This was achieved by modeling the average stimulation-evoked response of each neuron as linear function of two parameters: movement speed and stimulation current amplitude (linear regression; see Methods). We assessed the quality of each linear model by calculating a *R*^2^ value per neuron. To determine the *relative* modulation of neural responses by behavioral factors (speed) versus current amplitude, we took the absolute value of ratio of the two weights: 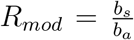. Taking the absolute value ignores whether neurons are inhibited or excited by each factor, and isolates only the magnitude of the contribution.

We found that neurons whose responses were better fit by linear regression (those with high *R*^2^ values) tended to weigh locomotion speed and stimulation amplitude equally (fig. 6c). In contrast, neurons will low *R*^2^ values placed a relatively lower weight on stimulation current amplitude, possibly indicating that they were unresponsive or that their responses were nonlinearly related to stimulation amlitude. Of note is that approximately ten percent of neurons had ratios below one, a level at which movement speed was the dominant driver of neural activity. Neural modulation by locomotion and electrical stimulation could be either excitatory or inhibitory, as gauged from the weight assigned in linear regression to each factor. We found that twenty-two percent of neurons (1513 neurons) were inhibited by locomotion while eighteen percent of neurons (1207 neurons) were inhibited by electrical stimulation. Approximately five percent of all neurons were inhibited by both locomotion and electrical stimulation (321 neurons). Together, these data show that behavioral factors such as movement speed can significantly affect stimulation-evoked responses.

## 4 Discussion

### Novel findings

We identified two prominent patterns of neural activation with current amplitude: 1. *spreading out*, where increasing stimulus amplitude recruits neurons located further from the electrode tip, and 2. *filling in*, where increasing stimulus amplitude would make neurons at a fixed distance more likely to be modulated by stimulation. The temporal response profile of neurons at low stimulation amplitudes was dominated by low-latency excitation. However, increasing current amplitude shifted response profiles in a distance-dependent manner. Within 300 *µ*m of the electrode tip, high stimulation amplitudes drove neurons to have long-latency, high-amplitude responses. In contrast, at longer distances increasing current amplitude yielded a growing fraction of inhibitory responses. When we examined only newly-activated neurons at each stimulus amplitude (those that had not been previously modulated by stimulation), we found more consistent responses across stimulation amplitudes: most neurons had low-latency excitatory responses, although a smaller fraction were inhibited. Despite following these general principles, the modulation of excitatory and inhibitory neurons by electrical stimulation is significantly different, as described below.

### High stimulation amplitudes biases network towards inhibition

The fraction of excitatory and inhibitory neurons activated by electrical stimulation scales will stimulation current amplitude at the same rate until 50 *µ*A, where inhibition fractionally but significantly dominates, reaching a ratio of 1.1 (fig. 2b). This is clearly seen in Figure 4b-c, where neural response profiles that show inhibition relative to baseline become more prominent, an effect that is particularly prominent at distances longer than 500 *µ*m from the electrode tip. A second indication that inhibition grows with stimulation amplitude is found in neurons adjacent to the stimulating electrode, where neural excitatory responses shift towards longer latencies (fig. 4b-c), possibly due to increasing recruitment of local inhibitory neurons.

### Lateral spread of excitation is impeded by distant inhibitory activation

The analyses describing the spatial spread of stimulation-evoked neural activity in Figures 3 and 4 separated inhibitory from excitatory neurons. From these datasets, it is clear that the lateral spread of inhibitory activity is faster than the lateral spread of excitatory activity at each stimulation amplitude. This observation holds across specific temporal response profiles, but is particular evident for both short-latency and long-latency activation of neurons (fig. 4b). In responses to sensory or artificial stimulation, several studies examined how each subtype of interneurons population influence the spatio-temporal dynamics of nearby excitatory populations, revealing that some of their effects can propagate for hundreds of *µ*m [11]. Decoupling inhibitory neurons by knocking out connexin 36, a neuronal gap junction protein that allows fast signal transmission across neurons, results in prolonged excitatory responses, a larger spatial extent of stimulation-evoked activity, and strong high-frequency oscillations [2].

### Behavioral modulation of stimulation-evoked activity

The variability of neural responses to electrical stimulation across trials is striking (fig. 6), and the mean fluctuations in the population across time indicate some changes in internal states of the animal, which can shift the dynamics of inhibition to excitation and in turn change the way that neurons respond to incoming sensory signals [16, 13, 20, 18, 4]. The fluctuations in our data were reasonably well correlated with running speed of the mouse, but may instead reflect some other internal variable such as arousal that can vary independently of locomotion [1]. Of note, a prior study found that electrical stimulation delivered during rest evokes low-frequency oscillations that are quenched during active sensorimotor behavior (whisking), even when rats were actively engaged in a behavioral task requiring they detect electrical stimulation inputs [26]. These data suggest that electrical stimulation intended to modulate neural activity in a specific pattern must take into account the ongoing activity of the local neural population or, at a courser scale, make some inference by tracking metrics regarding behavior and arousal.

### Activation patterns across the neural population

Together, the observations described above lead to a simple description of changing activity patterns across the population as a function of increasing stimulation amplitude: neurons close to the electrode tip are most likely to be modulated by stimulation and show excitatory low-latency responses. Increasing current amplitude increases the spatial extent of neurons activated by stimulation, but neurons closest to the electrode tip move toward longer-latency responses. In addition, network activity that controls the lateral spread of stimulus-evoked activity involves widespread activation of inhibitory neurons, which serve to keep the stimulation-evoked responses from growing too large but fail to focus it closely around the point of stimulation. These insights will be useful in designing stimulation parameters for protocols intended to shift the balance of excitation to inhibition within a brain region.

## 5 Acknowledgements

The work completed in this article was supported by by NIH grant EY02874 and the UCSF Program in Breakthrough Biomedical Research.

